# Systematic Perturbation of Thousands of Retroviral LTRs in Mouse Embryos

**DOI:** 10.1101/2023.09.19.558531

**Authors:** Jian Yang, Lauryn Cook, Zhiyuan Chen

## Abstract

In mammals, many retrotransposons are de-repressed during zygotic genome activation (ZGA). However, their functions in early development remain elusive largely due to the challenge to simultaneously manipulate thousands of retrotransposon insertions in embryos. Here, we employed epigenome editing to perturb the long terminal repeat (LTR) MT2_Mm, a well-known ZGA and totipotency marker that exists in ∼2667 insertions throughout the mouse genome. CRISPRi robustly repressed 2485 (∼ 93%) MT2_Mm insertions and 1090 (∼55%) insertions of the closely related MT2C_Mm in 2-cell embryos. Remarkably, such perturbation caused down-regulation of hundreds of ZGA genes at the 2-cell stage and embryonic arrest mostly at the morula stage. Mechanistically, MT2_Mm/MT2C_Mm primarily served as alternative ZGA promoters activated by OBOX proteins. Thus, through unprecedented large-scale epigenome editing, we addressed to what extent MT2_Mm/MT2C_Mm regulates ZGA and preimplantation development. Our approach could be adapted to systematically perturb retrotransposons in other mammalian embryos as it doesn’t require transgenic animals.

## 1. Introduction

Retrotransposons comprise ∼40% of the human genome (Lander et al., 2001). Precise control of retrotransposon activity is central to genome integrity because ectopic de-repression of retrotransposons have been linked to aging and numerous diseases (Gorbunova et al., 2021). However, some retrotransposons have also co-opted regulatory activities to control host gene expression and biological processes (Fueyo et al., 2022). For example, retrotransposons have been reported to function as promoters, enhancers, and silencers and to regulate 3D genome organizations (Fueyo et al., 2022; Modzelewski et al., 2022). Thus, fully understanding how retrotransposons are involved in normal development and disease progressions is an important biological question.

Retrotransposons are generally silenced in somatic cells but some of them are highly expressed in germ cells and early embryos (Fueyo et al., 2022). It is believed that these retrotransposons act as *cis*-regulatory elements to enable successful gametogenesis and early development. For example, MTC- and MT2B2-derived alternative promoters have been shown to be essential for mouse oogenesis and preimplantation development, respectively (Flemr et al., 2013; Modzelewski et al., 2021). In these two studies, experiments were carried out by knocking out (KO) single MTC and MT2B2 insertions. In addition to single copy KO, small interfering RNA (siRNA), antisense oligos (ASO), TALEN- and CRISPR-based methods have been used to simultaneously target many retrotransposon insertions (Fuentes et al., 2018; Huang et al., 2017; Jachowicz et al., 2017; Percharde et al., 2018; Sakashita et al., 2023; Todd et al., 2019; Yu et al., 2022). One drawback of siRNA- and ASO-mediated perturbation is that on- and off-targets can only be computationally predicted. In contrast, CRISPR-based approach allows for more accurate determination of targeted repeats by analyzing dCas9-binding (Fuentes et al., 2018).

MT2_Mm is a LTR of the mouse endogenous retroviruses with leucine tRNA primer (MERVL) element (Ribet et al., 2008). Both MT2_Mm and MERVL have been used as ZGA and totipotency markers in mice since 2012 (Genet and Torres-Padilla, 2020; Macfarlan et al., 2012). However, to what extent MT2_Mm regulates totipotency, ZGA, and early development remains unaddressed. This is largely due to the technical challenge to perturb ∼2667 MT2_Mm insertions throughout the mouse genome. MT2_Mm have been reported to form LTR-chimeric transcripts (<100) in 2-cell embryos (Macfarlan et al., 2012; Modzelewski et al., 2021; Peaston et al., 2004), but whether this chimeric transcription is essential for early development remains unclear. Here we address these questions by developing a computational and experimental framework to systematically perturb MT2_Mm and the related repeats in embryos.

## 2. Results

### 2.1. Targeting thousands of retroviral LTRs by multiplexed epigenome editing

To perturb MT2_Mm by epigenome editing, six single guide RNAs (sgRNAs) were designed to target ∼94.5% (2521/2667) MT2_Mm insertions in the current mouse genome assembly (**Fig. S1a**). Notably, ∼91.5% MT2_Mm insertions are predicted to be targeted by at least two sgRNAs with zero mismatch, and ∼79.6% are targeted by at least four sgRNAs (**Fig. S1a**). This design is critical because multiple sgRNA targeting increases the likelihood of robust epigenome editing (Perez-Pinera et al., 2013). About 5.5% (146) MT2_Mm insertions are not recognized by any sgRNAs. This is likely because they are too truncated compared to the MT2_Mm consensus (**Fig. S1b**). Since MT2C_Mm consensus is highly similar to MT2_Mm (85% identity), sgRNAs were not penalized for off-targeting if they also target MT2C_Mm. As a result, ∼28% MT2C_Mm copies are recognized by the sgRNAs with zero mismatch (**Fig. S1c**). These sgRNAs are predicted to have minimal off-targeting to other MT2 subfamilies including MT2B, MT2B1, and MT2B2 (**Fig. S1c**).

We next evaluated sgRNA on- and off-target effects in mouse embryonic stem cells (ESCs), an *in vitro* approximate of early embryos. We multiplexed the six sgRNAs using the CARGO (Chimeric Array of gRNA Oligos) method (Gu et al., 2018) and generated a transgenic ESC line constitutively expressing these sgRNAs (**Fig. 1a, Fig. S1d and S1e**). The ESC line also harbors a Dox-inducible catalytically dead Cas9 fused to a KRAB domain and an HA-tag (dCas9-KRAB-HA) (**Fig. 1a, Fig. S1d and S1e**). After Dox induction, chromatin immunoprecipitation followed by sequencing (ChIP-seq) was performed using antibodies against dCas9 and HA. The parental ESC line with the inducible dCas9-KRAB-HA, but not the MT2_Mm sgRNAs, served as a control.

**Fig. 1.**
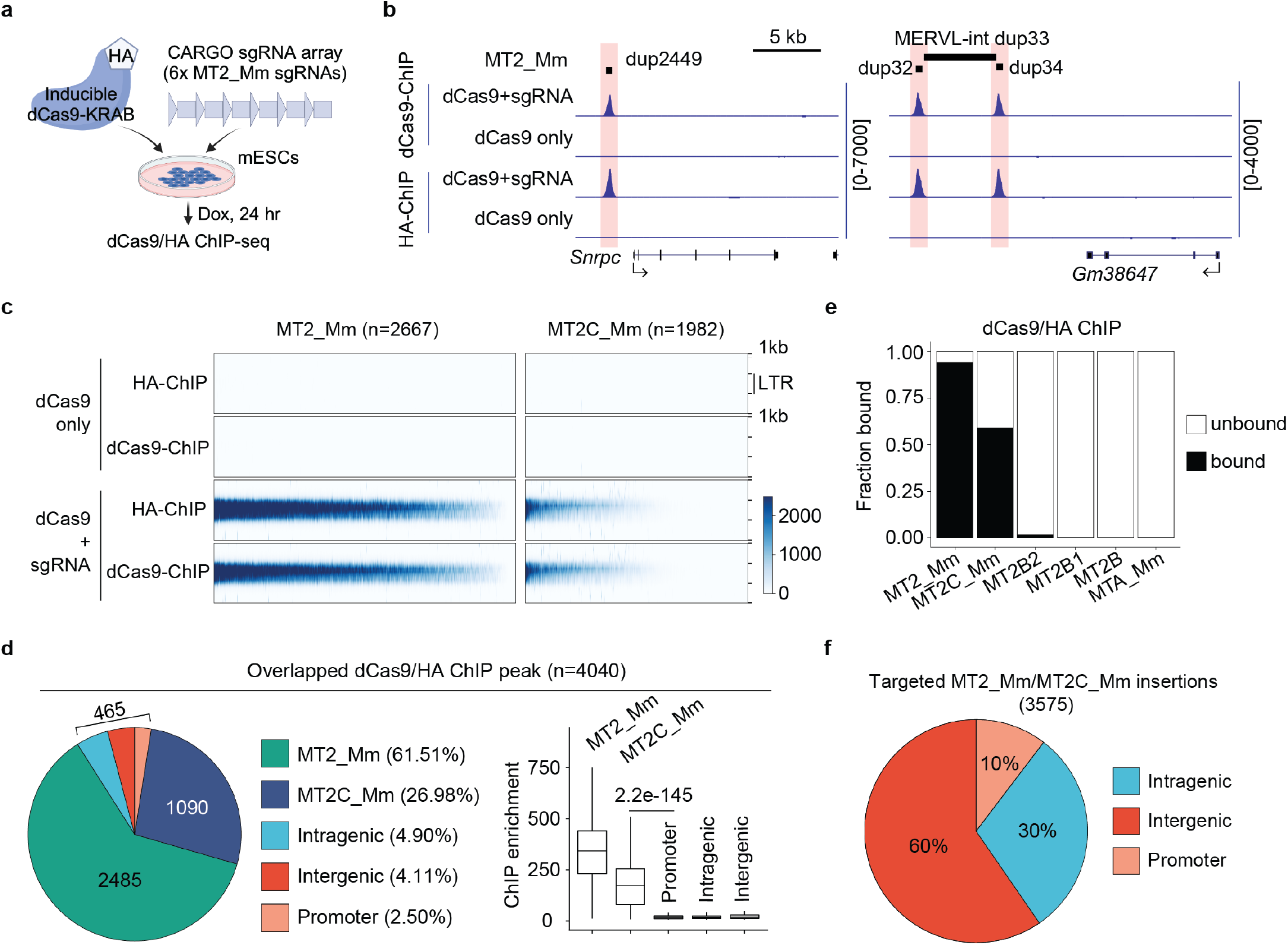
Selective dCas9 targeting to MT2_Mm/MT2C_Mm by CARGO. **a)** Experimental design. **b)** Genome browser views of dCas9 and HA ChIP signals at a solo MT2_Mm (left panel) and two MT2_Mm insertions flanking a full length MERVL (right panel). For ChIP-seq analyses, one alignment was randomly selected for read pairs that were mapped to multiple genomic locations to recover ChIP signals at repeats. **c)** Heatmap illustrating ChIP signals over all annotated MT2_Mm and MT2C_Mm insertions. **d)** Pie chart showing genomic distributions of high confidence ChIP-seq peaks (left panel). Box plot showing ChIP signals at the targeted MT2_Mm/MT2C_Mm insertions and the off-target sites (right panel). The middle lines in the boxes represent medians. Box hinges indicate the 25th and 75th percentiles, and whiskers indicate the hinge ±1.5 ×interquartile range. P-value, two-sided Wilcoxon rank-sum test. **e)** Stacked plot showing the fractions of LTR insertions bound by dCas9. **f)** Pie chart showing genomic distributions of the targeted MT2_Mm/MT2C_Mm insertions.

The dCas9 and HA ChIP peaks were highly overlapped at both MT2_Mm and MT2C_Mm insertions (**Fig. 1b and 1c**). We identified 4040 high confidence ChIP peaks (**Table S1**), with most of the dCas9-binding occurring at MT2_Mm (2485, 93.2% of MT2_Mm insertions) and MT2C_Mm sites (1090, 55.0% of MT2C_Mm insertions) (**Fig. 1d and 1e**). These targeted MT2_Mm/MT2C_Mm insertions were mostly localized in intergenic (60.0%) and intragenic regions (30.0%) (**Fig. 1f**). The remaining 465 peaks were classified as off-targets but had very low level of ChIP signals (**Fig. 1d**), suggesting weak and/or transient dCas9 binding. The off-target sites were considered in the downstream gene expression analyses. Lastly, no significant bindings were observed to other MT2 subfamilies (**Fig. 1e**). Thus, this data indicates that the CARGO-dCas9 selectively targets MT2_Mm and MT2C_Mm subfamilies.

### 2.2. Rapid, Robust, and specific CRISPRi in mouse embryos

To determine whether CRISPRi represses MT2_Mm expression in embryos, dCas9-KRAB mRNA and MT2_Mm sgRNAs were co-injected into 1-cell embryos (1C), and 2C embryos were collected for gene expression and immunostaining analyses (**referred to as MT2_Mmi**) (**Fig. 2a**). Both the non-injected embryos and the non-targeting sgRNA/dCas9-KRAB injection (**referred to as CTRi**) served as negative controls. Since MERVL-int is the internal viral element of MT2_Mm, the MERVL-int transcript and the encoded Gag protein were also assayed. Remarkably, both MT2_Mm and MERVL-int transcripts were efficiently repressed at the early 2C (E2C) stage (∼16hr post injection) (**Fig. S2a**). The ZGA genes *Zscan4* and *Obox3* were not affected by CTRi, suggesting minimal non-specific global transcriptional repression. In addition, the MERVL-Gag protein became undetectable in MT2_Mmi L2C embryos (**Fig. 2b and 2c**). The second polar bodies (arrows) served as internal immunostaining positive controls (**Fig. 2b**). It has been shown that the second polar body has both transcriptional and translational activities (Jin et al., 2022). The reason why MERVL expression in the polar body is not perturbed is likely due to that polar body has been extruded during micro-injection between 2-4 hpi (**Fig. 2a**). Note that depletion of the MERVL-Gag protein was achieved by injection of dCas9-KRAB mRNA at different concentrations.

**Fig. 2.**
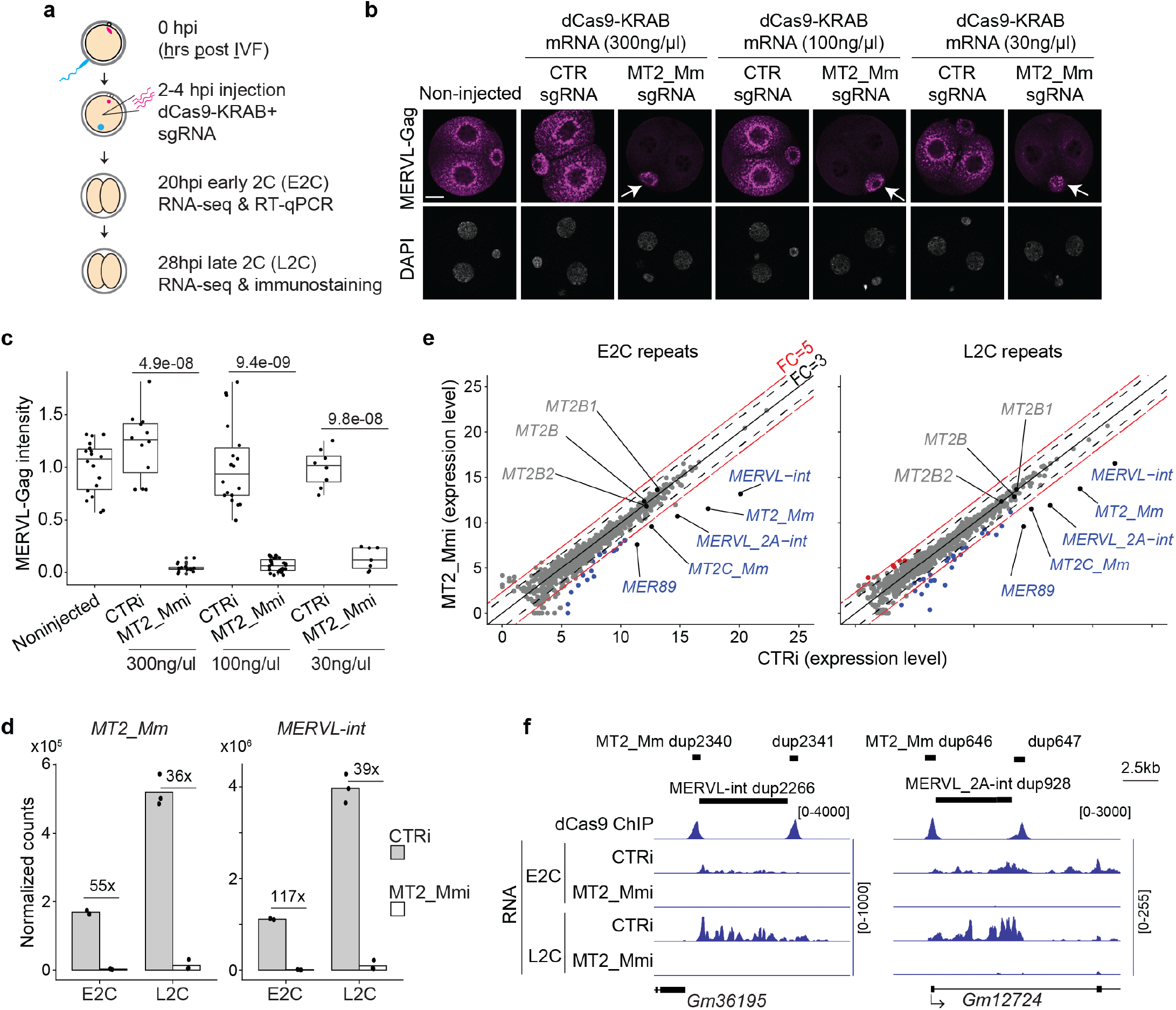
Rapid, robust, and specific perturbation of thousands of LTRs in embryos. **a)** Experimental design. IVF: *in vitro* fertilization. **b)** Immunostaining of MERVL-Gag protein at the L2C stage. White arrows point to the second polar body. Scale bar: 20µm. **c)** Quantifications of MERVL-Gag immunostaining signals. Each dot represents one 2C embryo. Number of embryos analyzed were 18, 12, 15, 18, 28, 8, and 7 for noninjected, CTRi (300ng/µl), MT2_Mmi (300ng/µl), CTRi (100ng/µl), MT2_Mmi (100ng/µl), CTRi (30ng/µl), and MT2_Mmi (30ng/µl), respectively. The middle lines in the boxes represent medians. Box hinges indicate the 25th and 75th percentiles, and whiskers indicate the hinge ±1.5× interquartile range. P-value, two-tailed t-test. **d)** Normalized RNA-seq read counts for MT2_Mm and MERVL-int at E2C and L2C stages. Both uniquely and multiple mapped read pairs were considered. Each dot represents one RNA-seq sample. **e)** Scatter plots comparing repeat expression at subfamily level between CTRi and MT2_Mmi. Differential repeat expression criteria are FC > 3 and FDR < 0.05. **f)** Genome browser views of dCas9 ChIP and RNA signals at a full length MERVL-int and a full length MERVL_2A-int. For RNA-seq tracks, only uniquely aligned read pairs are included.

Encouraged by these observations, total RNA-seq was performed to evaluate specificity and efficiency of CRISPRi in embryos. After confirming data reproducibility (**Fig. S2b**), we compared gene expression between CTRi and non-injected groups. Consistent with RT-qPCR, transcriptomes of these two groups were highly correlated (**Fig. S2c and S2d, Table S2**), indicating that CTRi did not have a global transcription repression effect. Notably, MT2_Mm and MERVL-int were down-regulated over 35-fold at both E2C and L2C stages (**Fig. 2d**). In addition, other related subfamilies including MT2C_Mm, MERVL_2A-int, and MER89 were among the top down-regulated repeats (**Fig. 2e and 2f, Fig. S2e, Table S2**). As expected, MT2B1, MT2B, and MT2B2 remained unchanged. Thus, this data demonstrated rapid (<16 hr), robust (>35-fold reduction), and specific CRISPRi-mediated perturbation of retrotransposons in embryos.

### 2.3. ZGA and preimplantation defects in MT2_Mmi embryos

We next sought to evaluate how MT2_Mmi may affect early development. Both CTRi and non-injected embryos cultured *ex vivo* developed to the blastocyst stage normally. However, most MT2_Mmi embryos were arrested at the morula stage (**Fig. 3a and 3b**). To determine how MT2_Mmi may cause the preim-plantation defect, we identified differentially expressed genes (DEGs) at E2C and L2C stages using a stringent cut off [fold change (FC) >3, false discovery rate (FDR) <0.05, and fragment per kilobase per million reads (FPKM) >1]. For both stages, more genes were down-regulated (206 for E2C and 471 for L2C) than up-regulated (52 for E2C and 91 for L2C) in the MT2_Mmi group (**Fig. S3a, Table S2**). This indicates that the predominant role of the targeted MT2_Mm/MT2C_Mm is to drive gene activation at these stages. Importantly, most down-regulated genes (90.3% for E2C and 82.4% for L2C) were within 50kb of the targeted repeats (**Fig. 3c**), suggesting that these repeats mainly activate genes *in cis*. In contrast, most up-regulated genes (84.6% for E2C and 91.2% for L2C) were >50kb away from the targeted repeats, suggesting an indirect effect. The down-regulated genes were enriched for gene ontology (GO) terms such as “ribonucleoprotein complex biogenesis” and “ribosome biogenesis” (**Fig. S3a**).

**Fig. 3.**
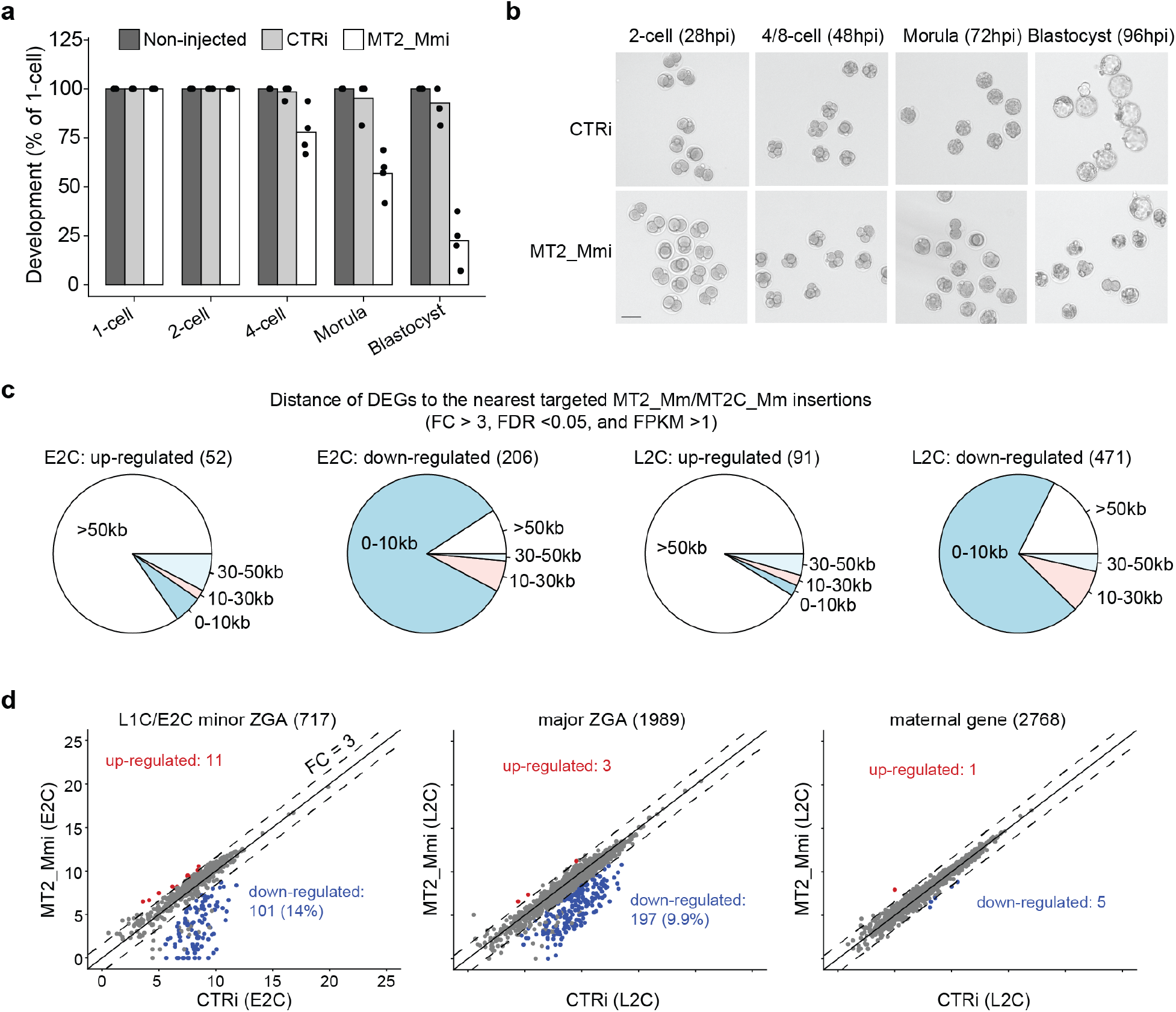
MT2_Mm/MT2C_Mm regulate ZGA and preimplantation development. **a)** Bar graph showing the percentage of embryos reaching 2C (28hpi), 4C (44hpi), morula (72hpi) and blastocyst (96hpi) stages. Dots represent the number of experiments performed. In total, 18, 38 and 52 embryos were analyzed for non-injected, CTRi, and MT2_Mmi groups, respectively. **b)** Bright field images showing embryos that reach to different developmental stages at indicated time points. Scale bar: 70µm **c)** Pie charts showing the distances of DEGs to the nearest targeted MT2_Mm/MT2C_Mm insertions. **d)** Scatter plots showing the expression level changes of minor ZGA, major ZGA, and maternal decay genes in the MT2_Mmi group.

We next evaluated how MT2_Mmi may affect ZGA. There are two waves of ZGA in mouse embryos. Minor ZGA takes place from L1C to E2C, whereas major ZGA occurs at L2C stage (Aoki et al., 1997; Bouniol et al., 1995; Schulz and Harrison, 2019). Using a previously reported cutoff (Ji et al., 2023), we identified 717 E2C minor ZGA genes, 1989 L2C major ZGA genes, and 2768 maternal genes (**Fig. S3b and Table S3**). A subset of minor (14%) and major ZGA genes (9.9%) were down-regulated at E2C and L2C, respectively (**Fig. 3d**). The ZGA defects should not be due to developmental delay because maternal decay at the L2C stage was not affected (**Fig. 3d**). Since MT2_Mmi embryos can develop to the morula stage (**Fig. 3b**), we also collected the healthiest 4C at 44 hpi and 8C at 54 hpi embryos for total RNA-seq analyses. The numbers of up- and down-regulated genes were comparable at these two stages (**Fig. S3c and Table S2**). The up-regulated genes at 4C stage were enriched for 2C genes (**Fig. S3d**), likely due to a delayed down-regulation. The GO terms enriched for the down-regulated genes at 4C and 8C included “blastocyst development” and “stem cell population maintenance” (**Fig. S3c**). These results suggest that MT2_Mmi-mediated ZGA defects at 2C manifest to cause gene dysregulation at later stages and underly the observed preimplantation phenotype.

### 2.4. MT2_Mm/MT2C_Mm primarily function as ZGA promoters activated by OBOX

To understand how MT2_Mmi caused gene dysregulation at 2C, we determined gene expression changes in relation to distances from the nearest targeted MT2_Mm/MT2C_Mm. We found that only host genes in direct proximity of the targeted repeats (<30 kb) were significantly down-regulated (**Fig. S4**). In contrast, such down-regulation was not observed for genes close to the off-targets identified by weak dCas9-binding (**Fig. 1d, and Fig. S4**). We further categorized the down-regulated genes at the 2C stages based on orientations and relative positions of host genes and the nearest targeted repeats (**Fig. 4a**). Remarkably, most of the down-regulated genes (92.3% for E2C and 83.3% for L2C) belonged to the “sense & upstream” category, that is, the nearest targeted MT2_Mm/MT2C_Mm was in sense direction and located upstream of the host gene (**Fig. 4b**). Given that enhancers activate gene transcription in an orientation- and position-independent manner, the strong bias toward “sense & upstream” category suggests that these repeats mainly function as promoters. In support of this, we identified MT2_Mm/MT2C_Mm-chimeric genes (175 for E2C and 207 for L2C) and found that these genes were significantly down-regulated upon MT2_Mmi (**Fig. 4c and Fig. 4g, Table S4**). However, a small subset of repeats may function as enhancers given that they overlapped with distal ATAC-seq peaks (**Fig. 4b and Fig. 4g**). Recently, oocyte-specific homeobox (OBOX) proteins were reported to regulate murine ZGA (Ji et al., 2023). We noted that OBOX activates a similar set of repeats to what we had successfully perturbed by epigenome editing (**Fig. 4d**). Since MT2_Mmi did not affect OBOX expressions (**Fig. 4e**), MT2_Mm/MT2C_Mm should act downstream of OBOX to activate the LTR-chimeric genes. We further identified 76 minor and 111 major ZGA genes that were down-regulated by both OBOX maternal-zygotic KO (mzKO) and MT2_Mmi (**Fig. 4f and 4g**). Thus, OBOX proteins regulate ZGA genes at least in part by activating MT2_Mm/MT2C_Mm-regulated genes.

**Fig. 4.**
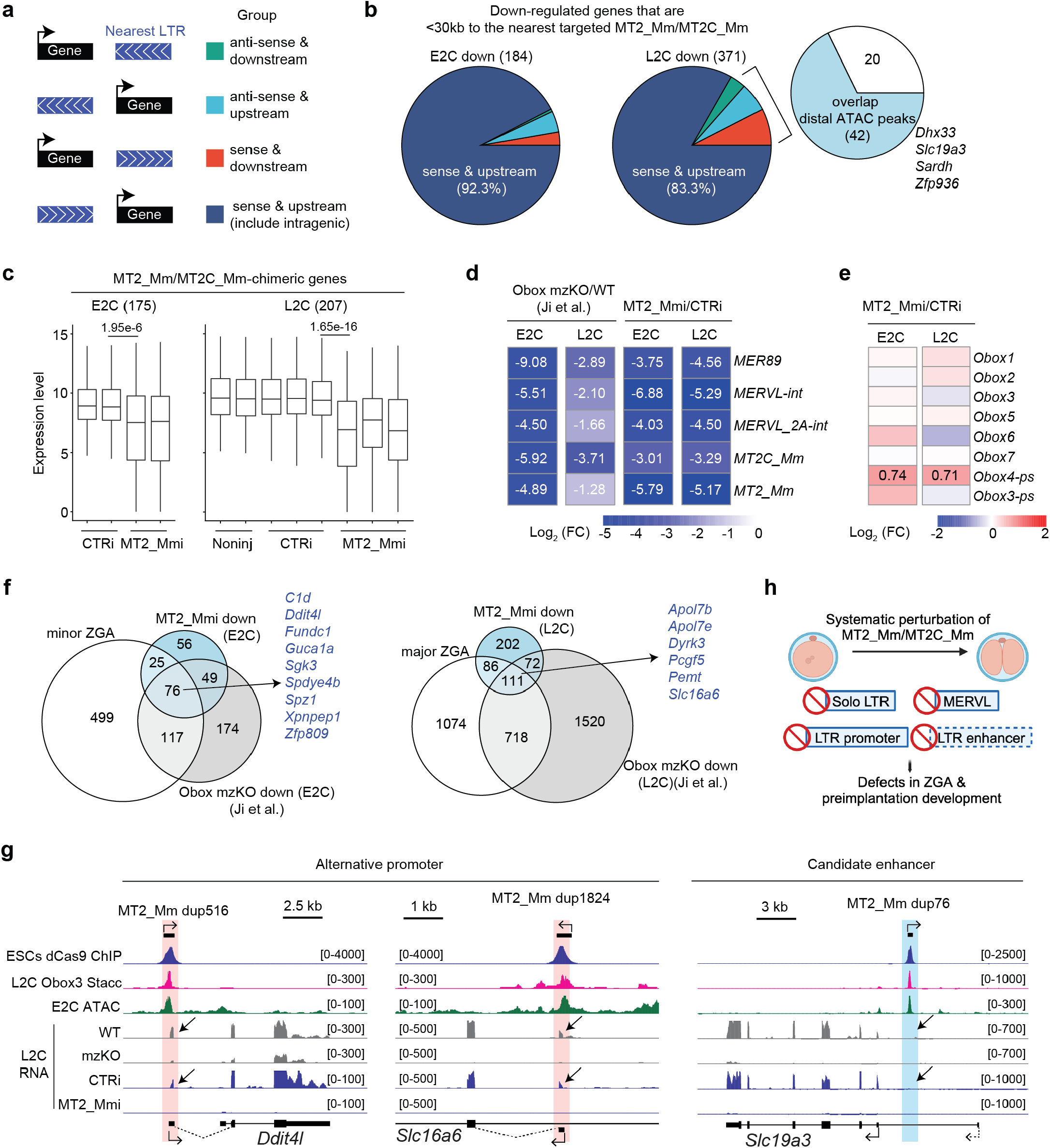
MT2_Mm/MT2C_Mm primarily serve as alternative promoters activated by Obox. **a)** Diagrams illustrating the possible relationships between genes and the nearest targeted MT2_Mm/MT2C_Mm insertions in terms of orientation and positions. Note that MT2_Mm/MT2C_Mm have their own orientations. **b)** Pie charts showing the relationships between down-regulated genes upon MT2_Mmi and the nearest targeted MT2_Mm/MT2C_Mm insertions. **c)** Box plots showing gene expression changes of MT2_Mm/MT2C_Mm-chimeric genes at E2C and L2C stages. The middle lines in the boxes represent medians. Box hinges indicate the 25th and 75th percentiles, and whiskers indicate the hinge± 1.5× interquartile range. P-value, two-sided Wilcoxon rank-sum test. **d)** Heatmap illustrating expression changes of the indicated repeat subfamilies in *Obox* maternal zygotic KO (mzKO) and MT2_Mmi groups. **e)** Heatmap illustrating expression changes of *Obox* genes upon MT2_Mmi. **f)** Venn diagrams showing the overlap between ZGA genes, and genes downregulated in both *Obox* mzKO and MT2_Mmi groups. **g)** Genome browser views of the indicated tracks at *Ddit4l, Slc16a6*, and *Slc19a3* locus. MT2_Mm orientations and gene transcription directions are as indicated. For *Ddit4l* and *Slc16a6*, 45-degree arrows point to the MT2_Mm-derived alternative promoters, and dashed lines indicate MT2_Mm-exon splicing junction. For *Slc19a3*, 45-degree arrow illustrates that no RNA signals are detected at the corresponding MT2_Mm insertion. **h)** Graphic summary of this work.

## 3. Discussion

Overall, our work achieved an unprecedented large-scale perturbation of retrotransposons in mouse embryos. We showed that MT2_Mmi perturbs both LTRs (i.e., MT2_Mm and MT2C_Mm) and their corresponding internal viral elements such as MERVL and MERVL_2A-int in embryos (**Fig. 4h**). MT2_Mmi also caused down-regulation of ZGA genes primarily by disrupting LTR-derived alternative promoters. Note that a small subset of MT2_Mm/MT2C_Mm may also function as enhancers. Collectively, these molecular alterations lead to preimplantation defects. To our knowledge, this is the first functional evidence of MT2_Mm/MT2C_Mm de-repression during ZGA and preimplantation development.

Recently, it was reported that MERVL promotes totipotency-to-pluripotency transition by repressing 2C genes through an unknown mechanism (Sakashita et al., 2023). This is an intriguing discovery given that MERVL is positively associated with 2C gene expression in 2-cell embryos, 2C-like cells, and totipotent-like cells (Nakatani and Torres-Padilla, 2023). In current study, de-repression of 2C genes was not observed despite that MERVL was strongly downregulated (>35-fold) upon MT2_Mmi (**Fig. 2 and 3**). In fact, MT2_Mmi mainly caused downregulation of the MT2_Mm/MT2C_Mm-chimeric ZGA genes at E2C and L2C stages (**Fig. 3 and 4**). The reason that causes the discrepancies remains unclear. One possibility is that ASOs used by Sakashita et a., only target MERVL, whereas our CRISPRi perturbs both MERVL and the flanking MT2_Mm/MT2C_Mm. Indeed, ASO-mediated knockdown of MERVL did not affect MT2_Mm-chimeric transcripts (Sakashita et al., 2023).

A few groups have employed epigenome editing to perturb retrotransposons in mouse embryos. Jachowicz et al. reported moderate reduction of LINE1 levels using a TALEN-based approach (Jachowicz et al., 2017). Sakashita et al. achieved ∼50% knockdown of MERVL mRNA by injecting a dCas9-KRAB-MECP2 plasmid into paternal pronuclei of zygotes (Sakashita et al., 2023). Here, we found that injection of dCas9-KRAB mRNA and targeting sgRNA can achieve more robust perturbation (>35 fold), which is likely due to that mRNA injection produces more abundant dCas9-KRAB proteins. In addition, cytoplasmic injection used in our method is technically simpler than pronuclei injection. Since our approach does not require transgenic animals, we expect that it could be adapted to investigate retrotransposon function at the subfamily level in other mammalian early embryos.

## 4. Materials and Methods

### 4.1. sgRNAs design and synthesis

MT2_Mm consensus sequence was retrieved from the Dfam database (Storer et al., 2021), and CRISPOR (Concordet and Haeussler, 2018) was used to design sgRNAs with the option “20bp-NGG-SpCas9”. The sgRNA target sites were predicted using Cas-OFFinder (Bae et al., 2014) allowing a maximum of three mismatches. The top six sgRNAs were picked based on the following criteria. First, sgRNAs were predicted to target most MT2_Mm insertions. Second, sgRNAs showed high on-target scores (i.e., “Doench ‘14-Score” and “Moreno-Mateos-Score”). Third, sgRNAs had minimal off-targeting to other MT2 subfamilies including MT2B, MT2B1, and MT2B2. Given the consensus sequences are highly similar between MT2C_Mm and MT2_Mm, sgRNAs predicted to target MT2C_Mm were not considered as off-targeting. The sgRNAs used for embryo micro-injection experiments were synthesized by Synthego with the following chemical modifications “2’-O-Methyl at 3 first and last bases, and 3’ phosphorothioate bonds between first 3 and last 2 bases”. All sgRNA sequences are available in **Table S5**.

### 4.2. dCas9-KRAB mRNA synthesis

The dCas9-KRAB-NLS-NLS sequence was amplified from the plasmid PB-TRE-dCas9-KRAB-MeCP2 (Tan, 2022) (addgene 122267) using PrimeSTAR Max 2 ×premix (Takara). A T7 promoter sequence was included in forward primer for *in vitro* transcription using the mMESSAGE mMACHINE T7 Ultra Transcription kit (Thermo Fisher Scientific). After LiCl precipitation, mRNA concentration was determined using a NanoDrop spectrophotometer, and stored at -80°C. Primer sequences are available in **Table S5**.

### 4.3. Generation of transgenic ESC lines

Mouse ESCs (E14TG2a, ATCC) were cultured on feeder-free dishes pre-coated with 0.1% gelatin (Sigma-Aldrich). ESCs were cultured in DMEM (high glucose, no glutamine, Thermo Fisher Scientific) medium supplemented with 15% Fetal Bovine Serum, 100U/ml Pen-streptomycin, 2mM GlutaMax, 1mM sodium pyruvate, 0.1mM non-essential amino acids, 0.084mM 2-mercaptoethanol, 1000 units/ml LIF, 0.5µM PD0325901, and 3µM ChIR99021. Cells were passaged 1:10-20.

To generate ESC lines carrying Dox-inducible dCas9, plasmid pT7077 (Hazelbaker et al., 2020) (addgene 137879) was co-transfected with Super piggyBac Transposase expression vector (System Biosciences) using Lipofectamine™ 3000 Transfection Reagent (Thermo Fisher Scientific). After transfection, cells were selected under 300µg/ml G418 (Fisher Scientific) for two days. After all cells in non-transfected negative control had died, G418 was supplemented at 100µg/ml throughout the rest of the experiments to avoid transgene silencing. Single colonies were picked 4-6 days later, and positive clones were selected based on GFP signal at 24 hr post Dox (1µg/ml)(Sigma Aldrich) induction.

The six MT2_Mm sgRNAs were assembled using the CARGO method (Gu et al., 2018) and inserted into pPN458 (Hazelbaker et al., 2020) BamH-I site (addgene 137876). After sequencing the whole plasmid using Plasmidsaurus, pPN4586× -MT2_Mm was co-transfected with Super piggyBac Transposase expression vector (System Biosciences) into an ESC line with Dox-inducible dCas9. After transfection, cells were selected under 10µg/ml Blasticidin S HCl (Thermo Fisher Scientific) for two days. Positive clones were picked based on RFP signal. GFP, RFP, and bright field images were acquired using EVOSII cell imaging system (Thermo Fisher Scientific).

### 4.4. Animal maintenance

Wild type B6D2F1/J (BDF1) (6- to 9-week-old) were purchased from Jax and maintained in the animal facility at Cincinnati Children’s Hospital Medical Center (CCHMC). All animal experiments were performed in accordance with the protocols of the Institutional Animal Care and Use Committee (IACUC) of CCHMC.

### 4.5. Collection of mouse embryos and micro-injection

Superovulation, metaphase II (MII) eggs collection, *in vitro* fertilization (IVF), embryo micro-injection, and embryo culture were described previously (Chen and Zhang, 2019). The time when sperm were added to cumulus oocyte complexes (COCs) was considered as 0 hr post IVF (hpi). About 6 pl dCas9-KRAB mRNA and sgRNA mixes were injected into cytoplasm of zygotes between 2-4 hpi using a Piezo impact-driven micromanipulator (Eppendorf). For all micro-injection experiments, sgRNAs were injected at 60ng/µl: 10ng/µl for each MT2_Mm sgRNA or 60ng/µl for the non-targeting control sgRNA. For micro-injection followed by immunostaining, dCas9-KRAB mRNA was injected at 300, 100, or 30ng/µl. For embryos used for analyses of developmental phenotypes, RTqPCR, and total RNA-seq, dCas9-KRAB mRNA was injected at 100ng/µl. Embryos were cultured in KSOM (Millipore) at 37°C under 5%CO2 with air. Embryo bright field images were acquired using EVOSII cell imaging system (Thermo Fisher Scientific).

### 4.6. Whole-mount immunostaining

Immunostaining was described previously (Inoue et al., 2018). The primary antibody was rabbit anti-MERVL-Gag (1:200, HUABIO, ER50102). The secondary antibody was Alexa Fluor 568 donkey anti-rabbit IgG (1:200) (Thermo Fisher Scientific). Fluorescence was detected under a laser-scanning confocal microscope (Nikon A1R inverted), and images were analyzed using ImageJ (NIH).

### 4.7. RT-qPCR

Twenty E2C (20hpi) embryos were collected and used for cDNA synthesis using SuperScript™ IV CellsDirect™ Lysis Reagents (Thermo Fisher Scientific). MT2_Mm, MERVL-int, *Obox3*, and *Zscan4* transcript abundance were determined using SYBR green gene expression assay (Thermo Fisher Scientific) in a QuantStudio 6 Flex system (Thermo Fisher Scientific). The Ct cycles were normalized to the *Actb* gene. The primers used are included in **Table S5**.

### 4.8. ChIP-seq library preparation and sequencing

About 10 million ESCs at 24 hr post Dox (1µg/ml) induction were collected for ChIP-seq library preparation. Cells were fixed in culture medium at room temperature for 10 min with final 1% formaldehyde (Sigma Aldrich), and fixation was quenched by final 0.14M glycine (Sigma Aldrich) at room temperature for 5 min. After cells were washed twice with icecold PBS (Thermo Fisher Scientific), cells were collected with scrapers. Following centrifugation (500g, 5 min, and 4°C) and the removal of supernatants, cell pellets were suspended in 1ml SDS lysis buffer (0.5% SDS, 100mM NaCl, 50mM Tris HCl-pH8.0, 5mM EDTA-pH8.0, 1× cOmplete EDTA free proteinase inhibitor cocktail), and incubated on ice for 10 min with a slight vortex every 2 min. After centrifugation (1000g, 5min, and room temperature), cell pellets were resuspended in 0.5ml IP buffer (0.3% SDS, 100mM NaCl, 50mM Tris HCl-pH8.0, 5mM EDTA-pH8.0, 1.6% Triton X-100, 1 ×cOmplete EDTA free proteinase inhibitor cocktail). Sonication was performed using Fisher Scientific Sonic Dismembrator with the following settings: 30% amplitude, ON=10s, OFF=30s for 8 cycles. Following sonication, supernatants containing chromatin were collected after centrifugation (20,000g, 4°C, 10 min). To determine chromatin concentration, about 10 µl chromatin was diluted using 190 µl elution buffer (1% SDS, 50mM Tris HCl-pH8.0, 1mM EDTA-pH8.0). Rest chromatin solution was aliquoted and stored at -80°C. Following reverse crosslinking (65°C, 600 rpm on a thermomixer, overnight), RNase A treatment (0.2mg/ml, 37°C, 1 hr), and Proteinase K (0.2mg/ml, 55°C, 1hr), DNA was isolated using UltraPure Phenol:Chloroform:Isoamyl Alcohol (Thermo Fisher Scientific) and quantified by a NanoDrop spectrophotometer. Chromatin containing 25µg DNA was used for each ChIP.

For each ChIP, 30µl protein A magnetic dynabeads (Thermo Scientific) were washed with IP buffer 3 times and incubated with primary antibody by rotation at 4°C for 4 hr. Primary antibodies were rabbit anti-Cas9 (Diagenode C15310258, 1:100) and rabbit anti-HA (Cell Signaling 3724S, 1:50). After preincubation, the beads/antibody were suspended and incubated with 100µl chromatin containing 25µg DNA at 4°C overnight on a rotator. The following day, beads/chromatin were washed once with low salt buffer (0.1% SDS, 150mM NaCl, 20mM Tris-HCl-pH8.0, 2mM EDTA, 1% Triton X-100), twice with high salt buffer (0.1% SDS, 500mM NaCl, 20mM Tris-HCl-pH8.0, 2mM EDTA-pH8.0, 1% Triton X-100), and twice with TE buffer (10mM Tris-HCl-pH8.0, 1mM EDTA-pH8.0). Chromatin was eluted by incubation with 200µl elution buffer at room temperature for 15 min and 37°C for 10 min on a 500rpm thermomixer. Eluted chromatin was reverse-crosslinked and treated with RNaseA and Proteinase K as described above. DNA was purified and quantified by Qubit (Thermo Fisher Scientific). About 1ng DNA was used for library construction using NEBNext Ultra II DNA Library Prep Kit for Illumina (NEB). Sequencing (2× 150bp) was performed on Illumina NovaSeq 6000 platform at Novogene. Library information is available in **Table S5**.

### 4.9. RNA-seq library preparation and sequencing

The total RNA-seq libraries were constructed using the SMART-Seq Stranded Kit (Takara) as previously described (Wang et al., 2022). For each library, 1µl 1/10000 ERCC RNA spike-in Mix was included prior to RNA shearing (Thermo Fisher Scientific). Sequencing (2× 150bp) was performed on the Illumina NovaSeq 6000 platform at Novogene. Library information is available in **Table S5**.

### 4.10. ChIP-seq data processing

For ChIP-seq data, the raw read pairs were trimmed by Trimgalore (v0.6.6) (https://github.com/FelixKrueger/TrimGalore) with the option “–paired”, and aligned to mm10 by bowtie2 (v2.4.2)(Langmead and Salzberg, 2012) with the options “-p 6 –no-unal –no-mixed –no-discordant -I 0, -X 1000”. Under this setting, one alignment was randomly selected for read pairs that were aligned to multiple genomic locations. PCR duplicates were removed using Picard (v2.18.22)(http://broadinstitute.github.io/picard/), and Macs2 (v2.1.4)(Zhang et al., 2008) was used to call peaks with the option “-f BAMPE -g mm -B -q 0.0001 –nolambda –nomodel”. For each antibody (i.e., anti-Cas9 and anti-HA), only peaks that were present in the “dCas9+sgRNA” group, but not the “dCas9-only”, were kept for downstream analyses. Overlaps between dCas9-ChIP and HA-ChIP were identified using bedtools (v2.27.0)(Quinlan and Hall, 2010) intersect function and considered as high confidence dCas9 target sites (n=4040). Bigwig files were generated using deeptools (v2.0.0)(Ramirez et al., 2016) bamCoverage function with the options “–binSize 25 –normalizeUsing RPKM –scalFactor 1”.

The dCas9/HA ChIP peaks were intersected to mm10 repeats annotations to identify peaks that overlapped with MT2_Mm and MT2C_Mm insertions. The pre-generated transposable elements (TE) GTF file “mm10_rmsk_TE.gtf.gz” was retrieved from the TEtranscripts (Jin et al., 2015) FTP site. The peaks were annotated using “makeTxDbFromGFF” and “annotatePeak” function from the “ChIPpeakAnno”, “GenomicFeatures”, and “ChIPseeker” Bioconductor R packages (Lawrence et al., 2013; Yu et al., 2015; Zhu et al., 2010). The gene annotation file was downloaded from GENCODE M25 (https://www.gencodegenes.org). The deeptools (v2.0.0)(Ramirez et al., 2016) “compueMatrix” and “plotHeatmap” functions were used to generate heatmaps illustrating dCas9/HA enrichment at MT2_Mm and MT2C_Mm insertions.

### 4.11. RNA data processing

Raw RNA-seq read pairs were trimmed by Trimgalore (v0.6.6)(https://github.com/FelixKrueger/TrimGalore) with the options “–paired –clip_R2 5”, and aligned to mm10 by STAR (v2.7.9)(Dobin et al., 2013) with the options “– outFilterMultimapNmax 100 –winAnchorMultimapNmax 100 –outSAMattributes NH HI NM MD XS AS –alignIntronMax 1000000 –alignMatesGapMax 1000000 –limitBAMsortRAM 30000000000 –outFilterType BySJout”. Following STAR mapping, TEtranscripts (v2.2.3)(Jin et al., 2015) was used to count uniquely and multiple aligned read pairs for each repeat subfamily. For genes, only uniquely mapped read pairs were counted. Uniquely aligned read pairs were further identified by sambamba (v0.6.8)(Tarasov et al., 2015) with the option “-F mapping_quality ≥255”, and then supplied for StringTie (v2.1.7)(Pertea et al., 2015) to generate FPKM values for each gene. Bigwig files were generated using deeptools (v2.0.0)(Ramirez et al., 2016) bamCoverage function with the options “–binSize 25 –normalizeUsing RPKM –scalFactor 1”. RNA-seq data were also aligned to a custom dCas9-KRAB and ERCC reference by bowtie2 (v2.4.2)(Langmead and Salzberg, 2012) to identify read pairs from dCas9-KRAB mRNA and ERCC RNA mixes.

The DESeq2 (v1.38.3)(Love et al., 2014) was used for differential expression analyses between different groups. For comparison between the non-injected group and CTRi, library size factors were estimated using the “estimateSizeFactors” function with ERCC read counts as the “controlGenes”. This comparison is to ensure that global transcriptional level was not affected by overexpression of dCas9-KRAB mRNA alone. For other comparisons, default DESeq2 settings were used. In addition to “FDR < 0.05”, additional cutoffs (mean FPKM of either group > 1, and fold change > 3) were used to define DEGs. For repeat subfamilies, only FDR < 0.05 and fold change > 3 were required. GO terms were analyzed using “enrichGO” function from Bioconductor R package “clusterProfiler” (v4.6.2). Distances of DEGs to nearest dCas9 MT2_Mm/MT2C_Mm ChIP peaks were determined using bedtools (v2.27.0)(Quinlan and Hall, 2010) closest function.

### 4.12. Define ZGA and maternal decay genes

The cutoffs used to define minor/major ZGA genes and maternal decay genes were described previously(Ji et al., 2023). The total RNA-seq data of MII eggs, L1C, E2C, and L2C were from two previous reports (Wang et al., 2022; Zhang et al., 2022). The reason why gene number for each category was different from the previous report (Ji et al., 2023) could be due to different reference genomes (mm9 vs. mm10) and gene annotations (UCSC vs. Gencode) being used. In fact, the ratio of affected minor and major ZGA genes upon *Obox* mzKO were comparable between this study and the previous analyses (Ji et al., 2023).

### 4.13. MT2_Mm/MT2C_Mm-chimeric gene identification

The total RNA-seq data of MII eggs, L1C, L2C, 4C, morula, and blastocyst from the previous report (Zhang et al., 2022) were used to identify LTR-chimeric genes as previously described (Modzelewski et al., 2021). The same pipeline was also used to identify LTR-chimeric genes using poly-A RNAseq datasets (Deng et al., 2014; Xue et al., 2013). LTR-chimeric genes in which MT2_Mm/MT2C_Mm served as promoters were combined from the three datasets.

### 4.14. Visualization and statistical analysis

All statistical analyses and plotting were performed using the R programming language. All genomic browser tracks were viewed using the UCSC genome browser (Kent et al., 2002).

## 5. Data availability

All sequencing data were deposited in the Gene Expression Omnibus under accession number GSE242123 (token for reviewers: xxxxxxxxx)

## 6. Code availability

The code for this study is available at https://github.com/ZYChen-lab/MT2_Mm-CRISPRi/.

## 7. Author contributions

Z.C. conceived and designed the experiments. Z.C., J.Y. and L.C. performed experiments. Z.C. analyzed the data. Z.C. wrote the manuscript with input from J.Y. and L.C.

## 8. Acknowledgements

We thank Drs. Yang Yu and Yueh-Chiang Hu for helping with mouse IVF and micro-injection system. We thank Drs. Azusa Inoue, Matthew T. Weirauch for the hepful discussions. This project was supported by Z.C.’s start-up funding from CCHMC. Z.C. is also supported by the Eunice Kennedy Shriver National Institute of Child Health and Human Development (R00HD104902).

## 9. Author information

The authors declare no competing financial interests.

---

**Fig. S 1.**
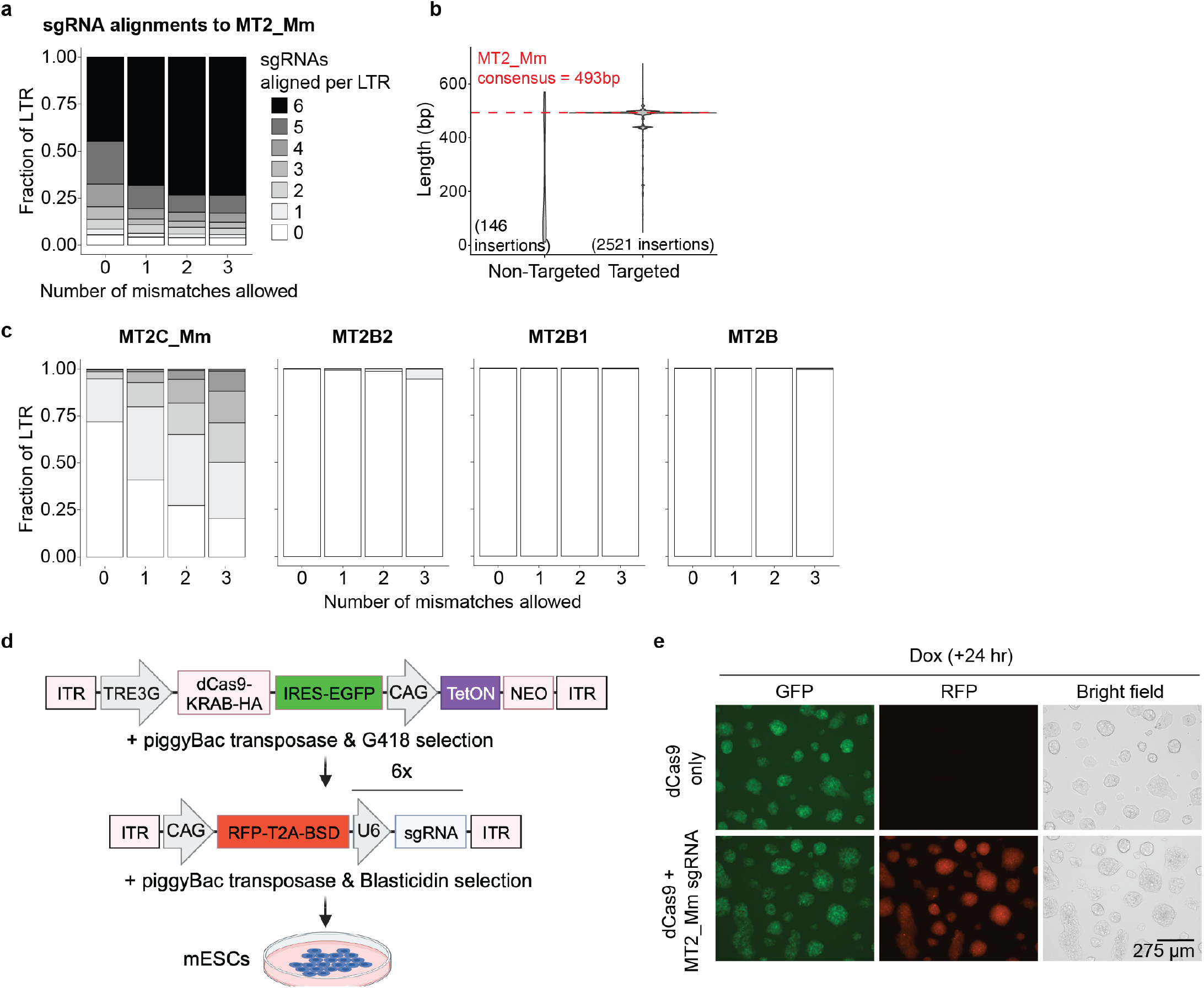
MT2_Mm sgRNA design and generation of ESCs for dCas9-binding analyses. **a)** Stacked plot showing the fractions of MT2_Mm insertions predicted to be targeted by different number of sgRNAs, with 0-3bp mismatches as indicated. White box depicts that 0 sgRNA is aligned, whereas black box indicates that all six sgRNAs are aligned. **b)** Violin plot showing the length distributions of MT2_Mm insertions that are predicted to be targeted by 0 sgRNA (non-targeted) or by at least one sgRNA (targeted). **c)** The same as panel a) except that different LTRs are presented. **d)** Diagrams illustrating the steps to generate ESCs carrying Dox-inducible dCas9 and MT2_Mm sgRNA array. **e)** GFP, RFP, and bright field images of the indicate ESC lines after Dox induction for 24 hr.

**Fig. S 2.**
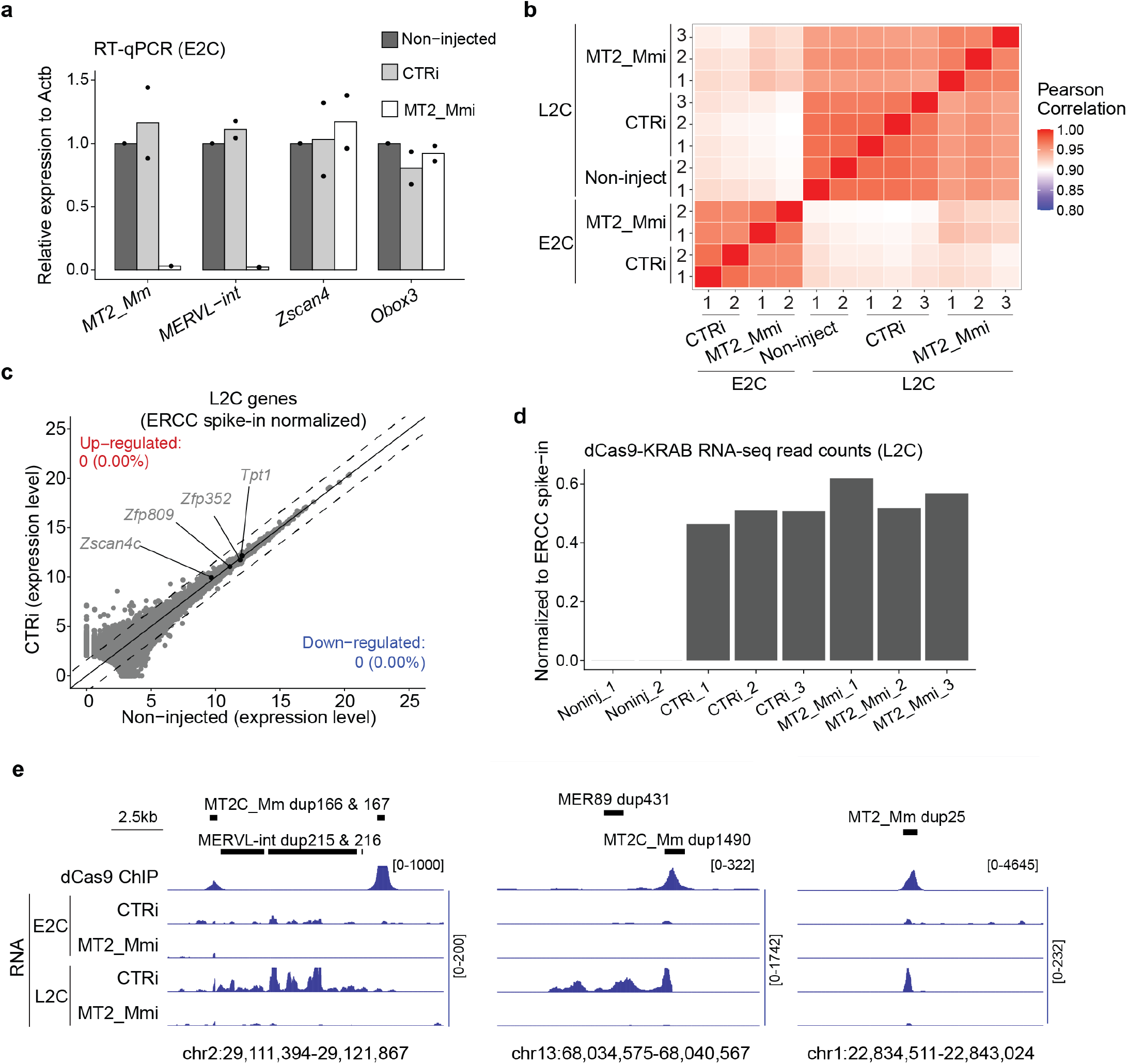
Quality control for 2C RNA-seq and CRISPRi perturbation. **a)** RT-qPCR analyses of the indicated genes and repeats at E2C stage. Each dot represents one biological replicate. **b)** Heatmap showing Pearson correlation between RNA-seq replicates. **c)** Scatterplot comparing gene expression changes between non-injected and CTRi groups. **d)** Bar graph showing the dCas9-KRAB RNA-seq read counts in the samples analyzed. **e)** Genome browser views of dCas9 ChIP and RNA signals at a full length MERVL-int (driven by MT2C_Mm), a MER89 (driven by MT2C_Mm), and a solo MT2_Mm. For RNA-seq tracks, only uniquely aligned read pairs are included.

**Fig. S 3.**
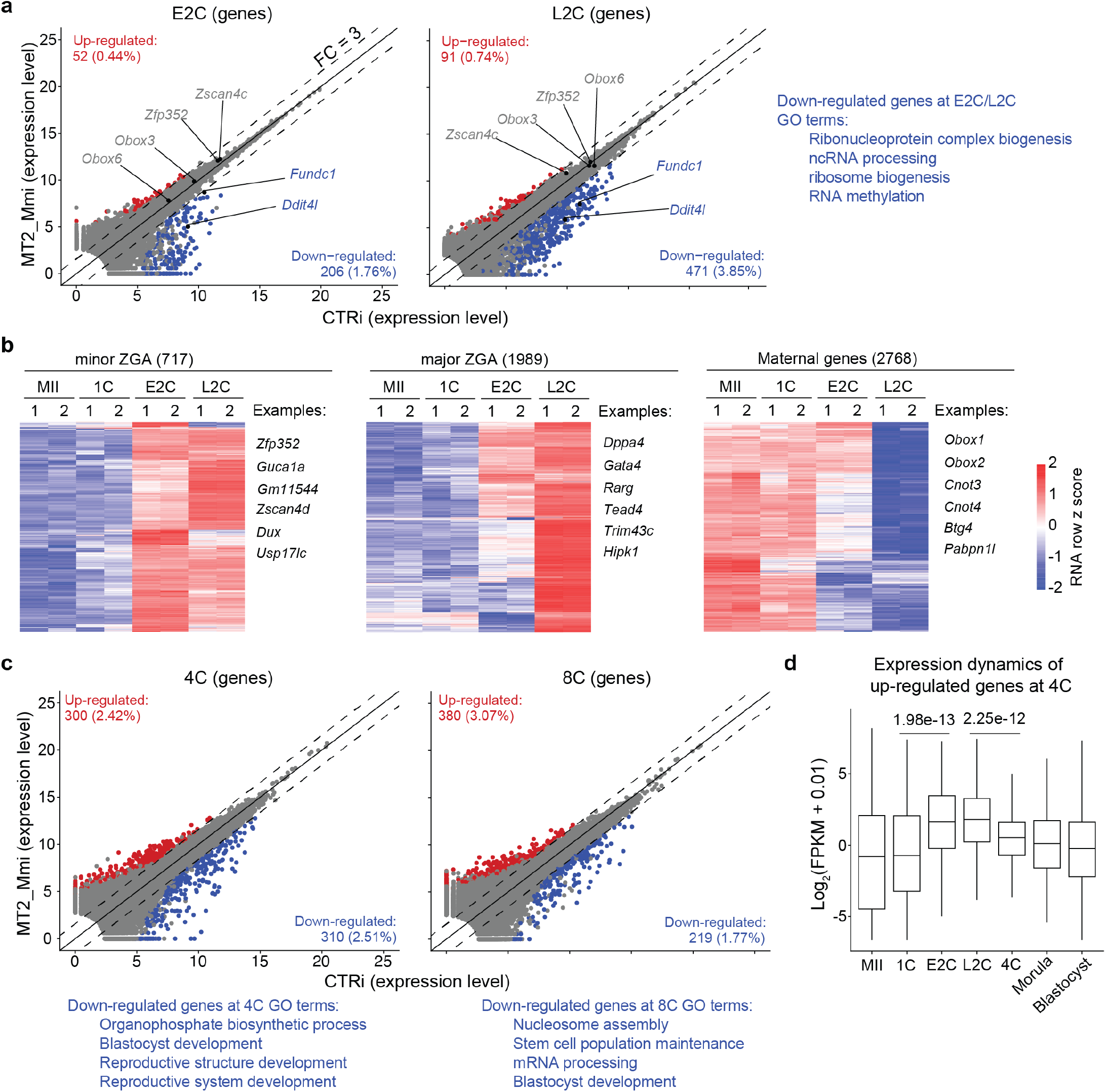
Impact of MT2_Mm/MT2C_Mm perturbation on early embryo transcriptome. **a)** Scatter plots comparing gene expression changes between CTRi and MT2_Mmi groups at E2C and L2C stages. Differential gene expression criteria are FC > 3, FDR < 0.05 and FPKM > 1. GO terms enriched are also listed. **b)** Heatmap illustrating the expression changes of minor ZGA, major ZGA, and maternal decay genes at the indicated stages. **c)** Scatter plots comparing gene expression changes between CTRi and MT2_Mmi groups at 4C and 8C stages. Differential gene expression criteria are FC > 3, FDR < 0.05 and FPKM > 1. GO terms enriched are also listed. **d)** Box plot showing the expression dynamics of up-regulated genes at 4C stage in the MT2_Mmi group. The middle lines in the boxes represent medians. Box hinges indicate the 25th and 75th percentiles, and whiskers indicate the hinge ±1.5× interquartile range. P-value, two-sided Wilcoxon rank-sum test.

**Fig. S 4.**
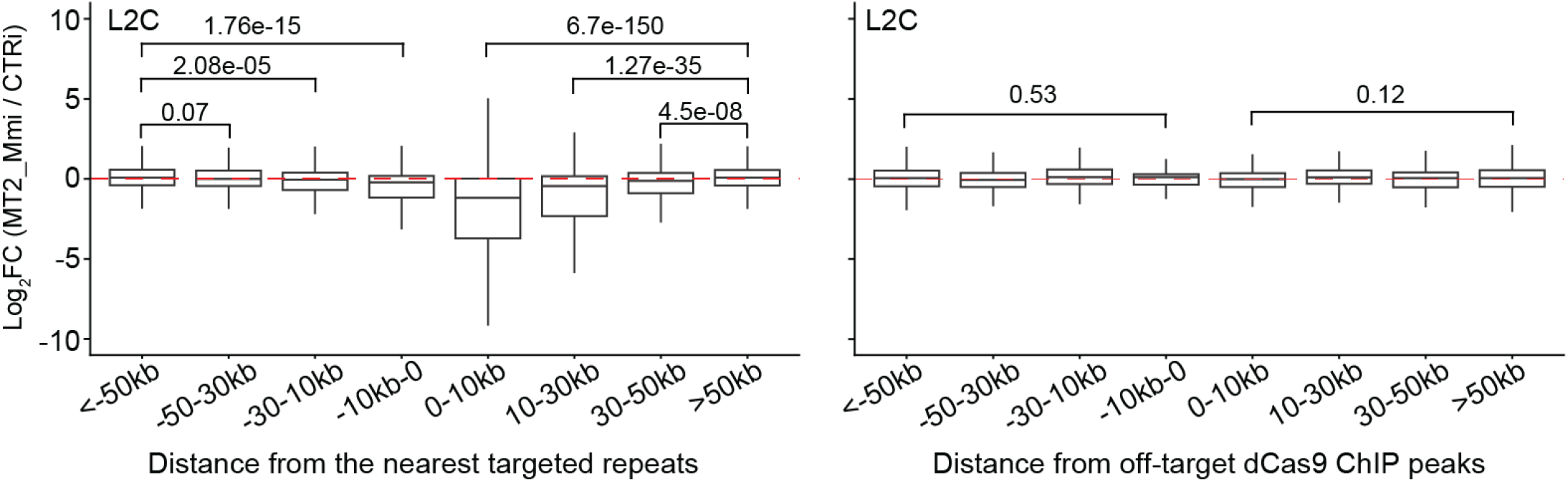
Gene expression changes in relation to distances from targeted repeats and off-target sites. The middle lines in the boxes represent medians. Box hinges indicate the 25th and 75th percentiles, and whiskers indicate the hinge±1.5×interquartile range. P-value, two-sided Wilcoxon rank-sum test.

## References

Aoki, F., Worrad, D.M., Schultz, R.M., 1997. Regulation of transcriptional activity during the first and second cell cycles in the preimplantation mouse embryo. Dev Biol 181, 296–307. doi:10.1006/dbio.1996.8466.

Bae, S., Park, J., Kim, J.S., 2014. Cas-offinder: a fast and versatile algorithm that searches for potential off-target sites of cas9 rna-guided endonucleases. Bioinformatics 30, 1473–5. doi:10.1093/bioinformatics/btu048.

Bouniol, C., Nguyen, E., Debey, P., 1995. Endogenous transcription occurs at the 1-cell stage in the mouse embryo. Exp Cell Res 218, 57–62. doi:10.1006/excr.1995.1130.

Chen, Z., Zhang, Y., 2019. Loss of dux causes minor defects in zygotic genome activation and is compatible with mouse development. Nat Genet 51, 947–951. doi:10.1038/s41588-019-0418-7.

Concordet, J.P., Haeussler, M., 2018. Crispor: intuitive guide selection for crispr/cas9 genome editing experiments and screens. Nucleic Acids Res 46, W242–W245. doi:10.1093/nar/gky354.

Deng, Q., Ramskold, D., Reinius, B., Sandberg, R., 2014. Single-cell rna-seq reveals dynamic, random monoallelic gene expression in mammalian cells. Science 343, 193–6. doi:10.1126/science.1245316.

Dobin, A., Davis, C.A., Schlesinger, F., Drenkow, J., Zaleski, C., Jha, S., Batut, P., Chaisson, M., Gingeras, T.R., 2013. Star: ultrafast universal rna-seq aligner. Bioinformatics 29, 15–21. doi:10.1093/bioinformatics/bts635.

Flemr, M., Malik, R., Franke, V., Nejepinska, J., Sedlacek, R., Vlahovicek, K., Svoboda, P., 2013. A retrotransposon-driven dicer isoform directs endogenous small interfering rna production in mouse oocytes. Cell 155, 807–16. doi:10.1016/j.cell.2013.10.001.

Fuentes, D.R., Swigut, T., Wysocka, J., 2018. Systematic perturbation of retroviral ltrs reveals widespread long-range effects on human gene regulation. Elife 7. doi:10.7554/eLife.35989.

Fueyo, R., Judd, J., Feschotte, C., Wysocka, J., 2022. Roles of transposable elements in the regulation of mammalian transcription. Nat Rev Mol Cell Biol 23, 481–497. doi:10.1038/s41580-022-00457-y.

Genet, M., Torres-Padilla, M.E., 2020. The molecular and cellular features of 2-cell-like cells: a reference guide. Development 147. doi:10.1242/dev.189688.

Gorbunova, V., Seluanov, A., Mita, P., McKerrow, W., Fenyo, D., Boeke, J.D., Linker, S.B., Gage, F.H., Kreiling, J.A., Petrashen, A.P., Woodham, T.A., Taylor, J.R., Helfand, S.L., Sedivy, J.M., 2021. The role of retrotransposable elements in ageing and age-associated diseases. Nature 596, 43–53. doi:10.1038/s41586-021-03542-y.

Gu, B., Swigut, T., Spencley, A., Bauer, M.R., Chung, M., Meyer, T., Wysocka, J., 2018. Transcription-coupled changes in nuclear mobility of mammalian cis-regulatory elements. Science 359, 1050–1055. doi:10.1126/science.aao3136.

Hazelbaker, D.Z., Beccard, A., Angelini, G., Mazzucato, P., Messana, A., Lam, D., Eggan, K., Barrett, L.E., 2020. A multiplexed grna piggybac transposon system facilitates efficient induction of crispri and crispra in human pluripotent stem cells. Sci Rep 10, 635. doi:10.1038/s41598-020-57500-1.

Huang, Y., Kim, J.K., Do, D.V., Lee, C., Penfold, C.A., Zylicz, J.J., Marioni, J.C., Hackett, J.A., Surani, M.A., 2017. Stella modulates transcriptional and endogenous retrovirus programs during maternal-to-zygotic transition. Elife 6. doi:10.7554/eLife.22345.

Inoue, A., Chen, Z., Yin, Q., Zhang, Y., 2018. Maternal eed knockout causes loss of h3k27me3 imprinting and random x inactivation in the extraembryonic cells. Genes Dev 32, 1525–1536. doi:10.1101/gad.318675.118.

Jachowicz, J.W., Bing, X., Pontabry, J., Boskovic, A., Rando, O.J., Torres-Padilla, M.E., 2017. Line-1 activation after fertilization regulates global chromatin accessibility in the early mouse embryo. Nat Genet 49, 1502–1510. doi:10.1038/ng.3945.

Ji, S., Chen, F., Stein, P., Wang, J., Zhou, Z., Wang, L., Zhao, Q., Lin, Z., Liu, B., Xu, K., Lai, F., Xiong, Z., Hu, X., Kong, T., Kong, F., Huang, B., Wang, Q., Xu, Q., Fan, Q., Liu, L., Williams, C.J., Schultz, R.M., Xie, W., 2023. Obox regulates mouse zygotic genome activation and early development. Nature doi:10.1038/s41586-023-06428-3.

Jin, H., Han, Y., Wang, H., Li, J.X.H., Shen, W., Zhang, L., Chen, L., Jia, S., Yuan, P., Chen, H., Meng, A., 2022. The second polar body contributes to the fate asymmetry in the mouse embryo. Natl Sci Rev 9, nwac003. doi:10.1093/nsr/nwac003.

Jin, Y., Tam, O.H., Paniagua, E., Hammell, M., 2015. Tetranscripts: a package for including transposable elements in differential expression analysis of rna-seq datasets. Bioinformatics 31, 3593–9. doi:10.1093/bioinformatics/btv422.

Kent, W.J., Sugnet, C.W., Furey, T.S., Roskin, K.M., Pringle, T.H., Zahler, A.M., Haussler, D., 2002. The human genome browser at ucsc. Genome Res 12, 996–1006. doi:10.1101/gr.229102.

Lander, E.S., Linton, L.M., Birren, B., Nusbaum, C., Zody, M.C., Baldwin, J., Devon, K., Dewar, K., Doyle, M., FitzHugh, W., Funke, R., Gage, D., Harris, K., Heaford, A., Howland, J., Kann, L., Lehoczky, J., LeVine, R., McEwan, P., McKernan, K., Meldrim, J., Mesirov, J.P., Miranda, C., Morris, W., Naylor, J., Raymond, C., Rosetti, M., Santos, R., Sheridan, A., Sougnez, C., Stange-Thomann, Y., Stojanovic, N., Subramanian, A., Wyman, D., Rogers, J., Sulston, J., Ainscough, R., Beck, S., Bentley, D., Burton, J., Clee, C., Carter, N., Coulson, A., Deadman, R., Deloukas, P., Dunham, A., Dunham, I., Durbin, R., French, L., Grafham, D., Gregory, S., Hubbard, T., Humphray, S., Hunt, A., Jones, M., Lloyd, C., McMurray, A., Matthews, L., Mercer, S., Milne, S., Mullikin, J.C., Mungall, A., Plumb, R., Ross, M., Shownkeen, R., Sims, S., Waterston, R.H., Wilson, R.K., Hillier, L.W., McPherson, J.D., Marra, M.A., Mardis, E.R., Fulton, L.A., Chinwalla, A.T., Pepin, K.H., Gish, W.R., Chissoe, S.L., Wendl, M.C., Delehaunty, K.D., Miner, T.L., Delehaunty, A., Kramer, J.B., Cook, L.L., Fulton, R.S., Johnson, D.L., Minx, P.J., Clifton, S.W., Hawkins, T., Branscomb, E., Predki, P., Richardson, P., Wenning, S., Slezak, T., Doggett, N., Cheng, J.F., Olsen, A., Lucas, S., Elkin, C., Uberbacher, E., Frazier, M., et al., 2001. Initial sequencing and analysis of the human genome. Nature 409, 860–921. doi:10.1038/35057062.

Langmead, B., Salzberg, S.L., 2012. Fast gapped-read alignment with bowtie 2. Nat Methods 9, 357–9. doi:10.1038/nmeth.1923.

Lawrence, M., Huber, W., Pages, H., Aboyoun, P., Carlson, M., Gentleman, R., Morgan, M.T., Carey, V.J., 2013. Software for computing and annotating genomic ranges. PLoS Comput Biol 9, e1003118. doi:10.1371/journal.pcbi.1003118.

Love, M.I., Huber, W., Anders, S., 2014. Moderated estimation of fold change and dispersion for rna-seq data with deseq2. Genome Biol 15, 550. doi:10.1186/s13059-014-0550-8.

Macfarlan, T.S., Gifford, W.D., Driscoll, S., Lettieri, K., Rowe, H.M., Bonanomi, D., Firth, A., Singer, O., Trono, D., Pfaff, S.L., 2012. Embryonic stem cell potency fluctuates with endogenous retrovirus activity. Nature 487, 57–63. doi:10.1038/nature11244.

Modzelewski, A.J., Gan Chong, J., Wang, T., He, L., 2022. Mammalian genome innovation through transposon domestication. Nat Cell Biol 24, 1332–1340. doi:10.1038/s41556-022-00970-4.

Modzelewski, A.J., Shao, W., Chen, J., Lee, A., Qi, X., Noon, M., Tjokro, K., Sales, G., Biton, A., Anand, A., Speed, T.P., Xuan, Z., Wang, T., Risso, D., He, L., 2021. A mouse-specific retrotransposon drives a conserved cdk2ap1 isoform essential for development. Cell 184, 5541–5558 e22. doi:10.1016/j.cell.2021.09.021.

Nakatani, T., Torres-Padilla, M.E., 2023. Regulation of mammalian totipotency: a molecular perspective from in vivo and in vitro studies. Curr Opin Genet Dev 81, 102083. doi:10.1016/j.gde.2023.102083.

Peaston, A.E., Evsikov, A.V., Graber, J.H., de Vries, W.N., Holbrook, A.E., Solter, D., Knowles, B.B., 2004. Retrotransposons regulate host genes in mouse oocytes and preimplantation embryos. Dev Cell 7, 597–606. doi:10.1016/j.devcel.2004.09.004.

Percharde, M., Lin, C.J., Yin, Y., Guan, J., Peixoto, G.A., Bulut-Karslioglu, A., Biechele, S., Huang, B., Shen, X., Ramalho-Santos, M., 2018. A line1-nucleolin partnership regulates early development and esc identity. Cell 174, 391–405 e19. doi:10.1016/j.cell.2018.05.043.

Perez-Pinera, P., Kocak, D.D., Vockley, C.M., Adler, A.F., Kabadi, A.M., Polstein, L.R., Thakore, P.I., Glass, K.A., Ousterout, D.G., Leong, K.W., Guilak, F., Crawford, G.E., Reddy, T.E., Gersbach, C.A., 2013. Rna-guided gene activation by crispr-cas9-based transcription factors. Nat Methods 10, 973–6. doi:10.1038/nmeth.2600.

Pertea, M., Pertea, G.M., Antonescu, C.M., Chang, T.C., Mendell, J.T., Salzberg, S.L., 2015. Stringtie enables improved reconstruction of a transcriptome from rna-seq reads. Nat Biotechnol 33, 290–5. doi:10.1038/nbt.3122.

Quinlan, A.R., Hall, I.M., 2010. Bedtools: a flexible suite of utilities for comparing genomic features. Bioinformatics 26, 841–2. doi:10.1093/bioinformatics/btq033.

Ramirez, F., Ryan, D.P., Gruning, B., Bhardwaj, V., Kilpert, F., Richter, A.S., Heyne, S., Dundar, F., Manke, T., 2016. deeptools2: a next generation web server for deep-sequencing data analysis. Nucleic Acids Res 44, W160–5. doi:10.1093/nar/gkw257.

Ribet, D., Louvet-Vallee, S., Harper, F., de Parseval, N., Dewannieux, M., Heidmann, O., Pierron, G., Maro, B., Heidmann, T., 2008. Murine endogenous retrovirus muerv-l is the progenitor of the “orphan” epsilon viruslike particles of the early mouse embryo. J Virol 82, 1622–5. doi:10.1128/JVI.02097-07.

Sakashita, A., Kitano, T., Ishizu, H., Guo, Y., Masuda, H., Ariura, M., Murano, K., Siomi, H., 2023. Transcription of mervl retrotransposons is required for preimplantation embryo development. Nat Genet 55, 484–495. doi:10.1038/s41588-023-01324-y.

Schulz, K.N., Harrison, M.M., 2019. Mechanisms regulating zygotic genome activation. Nat Rev Genet 20, 221–234. doi:10.1038/s41576-018-0087-x.

Storer, J., Hubley, R., Rosen, J., Wheeler, T.J., Smit, A.F., 2021. The dfam community resource of transposable element families, sequence models, and genome annotations. Mob DNA 12, 2. doi:10.1186/s13100-020-00230-y.

Tan, X; Worley, J.T.M.W.K.F.E.P.E.J.S.W.J.N.H.S.B.C.A.C.A., 2022. Interrogation of genome-wide, experimentally dissected gene regulatory networks reveals mechanisms underlying dynamic cellular state control. bioRxiv doi:10.1101/2021.06.28.449297.

Tarasov, A., Vilella, A.J., Cuppen, E., Nijman, I.J., Prins, P., 2015. Sambamba: fast processing of ngs alignment formats. Bioinformatics 31, 2032–4. doi:10.1093/bioinformatics/btv098.

Todd, C.D., Deniz, O., Taylor, D., Branco, M.R., 2019. Functional evaluation of transposable elements as enhancers in mouse embryonic and trophoblast stem cells. Elife 8. doi:10.7554/eLife.44344.

Wang, M., Chen, Z., Zhang, Y., 2022. Cbp/p300 and hdac activities regulate h3k27 acetylation dynamics and zygotic genome activation in mouse preimplantation embryos. EMBO J 41, e112012. doi:10.15252/embj.2022112012.

Xue, Z., Huang, K., Cai, C., Cai, L., Jiang, C.Y., Feng, Y., Liu, Z., Zeng, Q., Cheng, L., Sun, Y.E., Liu, J.Y., Horvath, S., Fan, G., 2013. Genetic programs in human and mouse early embryos revealed by single-cell rna sequencing. Nature 500, 593–7. doi:10.1038/nature12364.

Yu, G., Wang, L.G., He, Q.Y., 2015. Chipseeker: an r/bioconductor package for chip peak annotation, comparison and visualization. Bioinformatics 31, 2382–3. doi:10.1093/bioinformatics/btv145.

Yu, H., Chen, M., Hu, Y., Ou, S., Yu, X., Liang, S., Li, N., Yang, M., Kong, X., Sun, C., Jia, S., Zhang, Q., Liu, L., Hurst, L.D., Li, R., Wang, W., Wang, J., 2022. Dynamic reprogramming of h3k9me3 at hominoid-specific retro-transposons during human preimplantation development. Cell Stem Cell 29, 1031–1050 e12. doi:10.1016/j.stem.2022.06.006.

Zhang, C., Wang, M., Li, Y., Zhang, Y., 2022. Profiling and functional characterization of maternal mrna translation during mouse maternal-to-zygotic transition. Sci Adv 8, eabj3967. doi:10.1126/sciadv.abj3967.

Zhang, Y., Liu, T., Meyer, C.A., Eeckhoute, J., Johnson, D.S., Bernstein, B.E., Nusbaum, C., Myers, R.M., Brown, M., Li, W., Liu, X.S., 2008. Model-based analysis of chip-seq (macs). Genome Biol 9, R137. doi:10.1186/gb-2008-9-9-r137.

Zhu, L.J., Gazin, C., Lawson, N.D., Pages, H., Lin, S.M., Lapointe, D.S., Green, M.R., 2010. Chippeakanno: a bioconductor package to annotate chip-seq and chip-chip data. BMC Bioinformatics 11, 237. doi:10.1186/1471-2105-11-237.

